# Function-structure Coupling: White matter fMRI hyper-activation associates with structural integrity reductions in schizophrenia

**DOI:** 10.1101/2021.01.17.426982

**Authors:** Yuchao Jiang, Cheng Luo, Xiangkui Li, Huan Huang, Guocheng Zhao, Xuan Li, Shicai Li, Xufeng Song, Dezhong Yao, Mingjun Duan

## Abstract

**Background:** White matter (WM) microstructure deficit may be an underlying factor in the brain dysconnectivity hypothesis of schizophrenia using diffusion tensor imaging (DTI). However, WM dysfunction is unclear in schizophrenia. This study aimed to investigate the association between structural deficits and functional disturbances in major WM tracts in schizophrenia.

**Methods:** Using functional magnetic resonance imaging (fMRI) and DTI, we developed the skeleton-based white matter functional analysis, which could achieve voxel-wise function–structure coupling by projecting the fMRI signals onto a skeleton in WM. We measured the fractional anisotropy (FA) and WM low-frequency oscillation activation and their couplings in ninety-three schizophrenia patients and 122 healthy controls (HCs). An independent open database (62 schizophrenia patients and 71 HCs) was used to test the reproducibility. Finally, associations between WM activations and five behaviour assessment categories (cognition, emotion, motor, personality and sensory) were examined.

**Results:** This study revealed a reversed pattern of structure and function in frontotemporal tracts, as follows. (1) WM hyper-activation was associated with reduced FA in schizophrenia. (2) The function–structure association was positive in healthy controls but negative in schizophrenia patients. Furthermore, function–structure dissociation was exacerbated by long illness duration and severe negative symptoms. (3) WM activations were significantly related to cognition and emotion.

**Conclusions:** This study indicated function–structure dys-coupling, with higher functional activation and reduced structural integration in frontotemporal WM, which may reflect a potential mechanism in WM neuropathologic processing of schizophrenia.

## 1. Introduction

The mechanism of schizophrenia (SZ) has been postulated through the brain dysconnectivity hypothesis, positing that SZ is the result of ineffective or inefficient communications between brain regions (Dong, et al., 2019). As brain grey matter regions interconnect with white matter (WM), which is composed mainly of myelinated axons, WM damage may be an underlying factor in the pathological changes of SZ (Kelly, et al., 2018). Neuroimaging studies have also provided broad support for the disturbances of WM in both structural and functional organization in SZ (Dong, et al., 2017; Jiang, et al., 2019a). Furthermore, these abnormalities in WM have been linked to cognitive deficits, clinical symptoms and genetic alterations in SZ (Kochunov, et al., 2017; Monin, et al., 2015). Diffusion tensor imaging (DTI) has revealed widespread WM microstructure abnormalities in SZ patients (Kelly, et al., 2018). A recent meta-analysis consisting of 1963 SZ patients and 2359 healthy subjects revealed that WM microstructure alterations involve frontal, temporal and occipital lobes, as well as major fibre tracts connecting these regions, with lower fractional anisotropy (FA) in SZ patients (Kelly, et al., 2018).

As for WM function, increasing numbers of works have indicated the existence of brain functional activation in WM, as revealed by the blood oxygenation level-dependent (BOLD) signal in fMRI. For instance, specific regions in WM can be functionally activated in multiple tasks that correspond to different functional demands, including perceptual, language and motor processing demands (Courtemanche, et al., 2018; Gawryluk, et al., 2014a). Ji and colleagues found that the power of low-frequency BOLD fluctuations in the optic radiation was higher in the visual stimulation state than in the resting state, suggesting that BOLD signal in WM could be modulated by physiological states (Ji, et al., 2017). Additionally, BOLD signals within WM exhibited an intrinsic functional organization rather than random noise, even during the resting-state (Peer, et al., 2017). The anatomical bundles in WM can also be identified by DTI tracking with fMRI in the human brain (Ding, et al., 2013; Peer, et al., 2017). A recent study also reported the similarities between resting-state fMRI, DTI and histological connectivity in non-human primate brains (Wu, et al., 2019). This evidence indicates that the fMRI BOLD signal is a novel perspective for investigating WM dysfunction in brain disorders such as mild cognitive impairment, epilepsy, Alzheimer's and Parkinson's diseases (Ji, et al., 2019; Jiang, et al., 2019b; Makedonov, et al., 2016). More recently, our study revealed disrupted functional synchronization between WM and grey matter networks in SZ (Jiang, et al., 2019a; Jiang, et al., 2020).

However, it remains unclear whether structural and functional abnormalities detected by DTI and fMRI are localized in common WM tracts or whether there is an association between structural deficits and functional disturbances in WM-specific regions. To address these questions, we developed a DTI–fMRI fusion approach by projecting the WM microstructural features (FA) from DTI and the WM functional signals (BOLD time series) of fMRI onto a common WM skeleton based on the idea of tract-based spatial statistics (TBSS). Using this approach, we examined WM functions estimated by resting-state low-frequency oscillations in the WM skeleton in ninety-seven patients with SZ and 126 healthy subjects. We investigated common WM tracts with FA and WM function differences and further evaluated their associations.

## 2. Methods and Materials

### 2.1 Subjects

The current study recruited ninety-seven SZ patients and 126 healthy subjects (HC) from the Clinical Hospital of Chengdu Brain Science Institute (CHCBSI), China. Psychiatric symptom severity was assessed using the Positive and Negative Syndrome Scale (PANSS). History of major medical or neurological abnormalities, substance abuse, or other contraindications to MRI was considered as the exclusion criteria for all subjects. In addition, to eliminate potential familial effects, we further excluded healthy participants whose first- and second-degree relatives had a history of mental disorders. Two patients were first-episode and 95 were chronic. All of the SZ patients received antipsychotic medications. More information about the subjects is provided in Table 1. Written informed consent was signed by each participant before MRI scanning. The Ethics Committee of CHCBSI approved this study. Some of the patients were part of our previous studies and have been described in prior published studies (Dong, et al., 2019) (more details are listed in the Supplementary Materials).

**Table 1.** Demographic and clinical characteristics of subjects.

|  | Schizophrenia | Healthy controls | P value |
| --- | --- | --- | --- |
| Number | 93 | 122 | - |
| Gender (female/male) | 28/65 | 41/81 | 0.586<sup>a</sup> |
| Age (years) | 40.01±11.49 | 37.95±14.74 | 0.267<sup>b</sup> |
| Education (years) | 11.47±2.51 | 11.07±3.07 | 0.299<sup>b</sup> |
| White matter volume (mm<sup>3</sup>) | 489.96±57.46 | 495.65±55.09 | 0.463<sup>b</sup> |
| Disease duration (years) | 15.22±10.27 | - | - |
| Chlorpromazine equivalents (mg/day) | 337.96±145.26 | - | - |
| PANSS scores |  |  |  |
| Positive score | 13.32±5.89 | - | - |
| Negative score | 20.7±6.06 | - | - |
| General score | 28.19±5.86 | - | - |
| Total score | 62.21±13.26 | - | - |

<sup>||</sup> represents that four SZ patients and four HC were excluded from original database (97 SZ and 126 HC) because they failed to accomplish all of the T1, resting-state fMRI and DTI data acquisitions, or their head-motion beyond 2 mm or 2°.
<sup>a</sup> P-value was obtained by the Chi-square test
<sup>b</sup> P-value was obtained using the two-sample t-test

### 2.2 Data acquisition

High-resolution T1-weighted images, resting-state fMRI and diffusion-weighted images were acquired in a 3.0 Tesla GE MRI scanner (DISCOVERY MR 750, USA). Detailed scanning parameters are described in the Supplementary Material.

### 2.3 Data Processing: Skeleton-based White matter Functional Analysis

We developed a DTI–fMRI fusion approach by projecting the WM microstructural image (i.e., the FA image) and the WM functional images onto a common WM skeleton based on the idea of tract-based spatial statistics (TBSS). This method could achieve the assessment of function–structure coupling at voxel-wise in WM. The figure 1 provided a flowchart of the Skeleton-based White matter Functional Analysis (SWAF).

**Figure 1.**
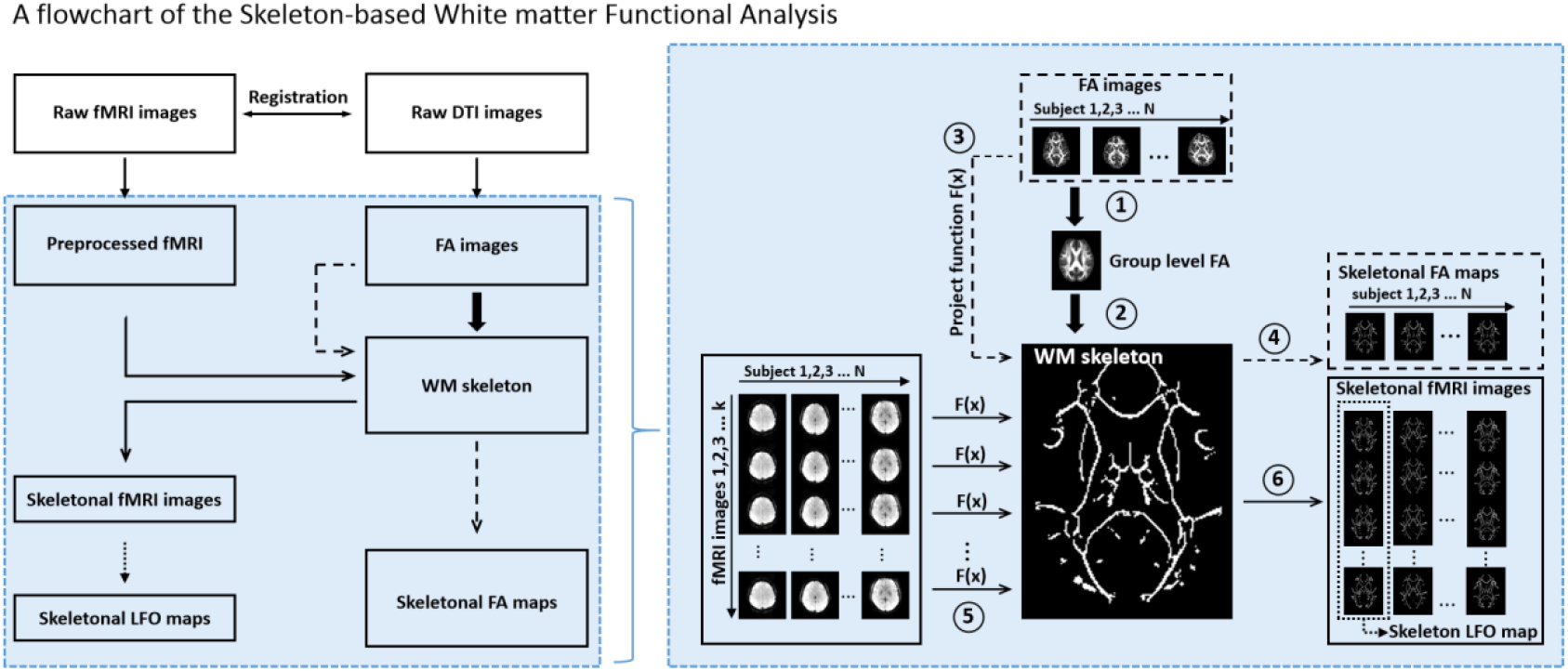
A flowchart of the Skeleton-based White matter Functional Analysis (SWAF), which was achieved by projecting the microstructural features (fractional anisotropy [FA]) of diffusion tensor imaging (DTI) and the white matter (WM) blood oxygenation level dependent (BOLD) signals of fMRI onto a common WM skeleton. (1) The raw DTI images were preprocessed using the FSL (https://www.fmrib.ox.ac.uk/fsl) and the individual FA image was further nonlinearly registered into a normalized MNI template. A group-level FA image was obtained by averaging across all subjects’ FA images. (2) The group-level FA image was thinned to create an WM FA skeleton. The WM skeleton represented the centres of all WM tracts common to the group. (3-4) Each subject’s FA image was projected onto the WM skeleton using the tract-based spatial statistics (TBSS) and yielded the projection function F(x). This projection generated a skeletonal FA map for each subject. (5-6) The raw fMRI images were registered into DTI b0 space and further preprocessed. Subsequently, the preprocessed fMRI images were projected onto the WM skeleton using the same projection function F(x) in the FA analysis. Finally, the skeletonal fMRI images were temporally filtered (band-pass, 0.01-0.08 Hz), and the amplitude of low frequency fluctuation was calculated to obtain the low-frequency oscillations (LFO) maps.

#### 2.3.1 DTI data processing

The DTI data were preprocessed in FSL (https://www.fmrib.ox.ac.uk/fsl). First, eddy current distortions, head motion correction and brain extraction were performed using FSL’s “eddy current correction” and the Brain Extraction Tool. Second, the FA image of each subject was estimated using DTIFIT (https://www.fmrib.ox.ac.uk/fsl/fdt/fdt_dtifit.html). Next, all subjects’ FA images were nonlinearly registered into a common space using the nonlinear registration tool FNIRT. Then, the mean FA images across all subjects was thinned to create an WM FA skeleton. The FA skeleton represented the centres of all WM tracts common to the group. Finally, each subject’s FA image was projected into the FA skeleton to obtain an individual FA skeleton map. In total, this analysis obtained all subjects’ FA skeleton maps, which were used for the following statistics.

#### 2.3.2 fMRI data processing

First, for each subject’s fMRI data, the first five volumes of fMRI time series were removed. Second, pre-processing procedures, including slice-timing correction and head motion correction, were performed. Subjects with maximum head motion > 2 mm or 2° were excluded. The mean frame displacement (FD) was also calculated to compare the difference between the two groups. Nuisance signals (24-parameter head motion and cerebrospinal fluid) and linear trend were further regressed out by a multiple linear regression model. Next, For each subject, the pre-processed functional images were registered to corresponding diffusion b0 image and then non-linearly projected onto the FA skeleton using the same transformation function in the DTI analysis. This resulted in a time series of skeletonized functional images. By projecting the functional images to the FA skeleton space, this procedure could theoretically improve sensitivity by reducing heterogeneity between individuals. In addition, this projection can reduce the methodological bias caused by misalignment and misregistration between DTI and fMRI space compared with that in conventional voxel-based analysis (Smith, et al., 2006).

#### 2.3.3 Low-frequency oscillations (LFO) of the WM skeleton

To characterize the WM activation, we estimate the LFO of the WM skeleton. The amplitude of low frequency fluctuation (ALFF) value (Zang, et al., 2007) was calculated for each voxel and yielded the skeleton-based WM amplitude of low frequency fluctuation (SWALFF) map. Specifically, the time series of a voxel were transformed into the frequency domain using the fast Fourier transform (FFT) to obtain the power spectrum. The square root was calculated at each frequency and the averaged square root across 0.01–0.1 Hz was taken as the ALFF value. For standardization purposes, the ALFF of each voxel was divided by the skeleton-averaged ALFF value for each subject (Yan, et al., 2013).

#### 2.3.4 Dynamic LFO of the WM skeleton

To further characterize the temporal dynamics of the LFO in the WM skeleton, we measured time-varying LFO using a sliding window approach (Dong, et al., 2019) by quantifying the temporal variability of the SWALFF. Previous studies indicated that window lengths less than 1/*f*_*min*_ may increase the risk of spurious fluctuations in the computation of dynamic LFO (Leonardi and Van De Ville, 2015). The *f*_*min*_ was the minimum frequency of the time series of resting-state fMRI. A longer sliding window length may hinder the characterization of the temporal variability in the dynamic LFO (Leonardi and Van De Ville, 2015). Thus, we used the optimal window length of 50 TRs (100 seconds) to calculate the dynamics of LFO. Using a window length of 50 TRs and a sliding step of five TRs, the full time series of 250 TRs (500 seconds) were divided into 40 windows for each subject. In each sliding window, the SWALFF map was calculated with a frequency band of 0.01–0.1 Hz. Subsequently, the coefficient of variation (CV) of SWALFF maps across sliding windows was computed to measure the dynamics of the SWALFF (dSWALFF).

### 2.4 Statistics: differences in FA, SWALFF and dSWALFF between SZ patients and HCs

For FA skeleton maps, SWALFF maps and dSWALFF maps, two-sample t-tests were used to compare the differences between SZ patients and HCs. Age, gender and education years were considered as covariates. Multiple comparisons correction was performed using the threshold-free cluster enhancement method (P<0.05, FWE correction) (Smith and Nichols, 2009).

To validate whether these WM differences between SZ patients and HCs from the current sample could also be observed on another independent SZ database, a publicly available SZ database (COBRE) (Cetin, et al., 2014), including raw anatomical, diffusion and fMRI data from 62 patients with SZ and 71 HCs, was used to calculate the FA, SWALFF and dSWALFF maps. The detailed subject information from the CBORE database is shown in the Supplementary Material.

### 2.5 Overlapping of dysfunctional WM tracts with microstructural deficits

To further investigate the associations between WM dysfunction and microstructural deficits, the tracts overlapping with FA-altered voxels and SWALFF-altered or dSWALFF-altered voxels were extracted. The Kolmogorov-Smirnov tests were used to examine the normal distribution of the variables. Subsequently, Pearson’s correlation coefficient was calculated between FA values and SWALFF or dSWALFF values in the overlapping tracts.

### 2.6 Relationships between WM function–structure coupling and clinical variables in SZ patients

We investigated the relationships between WM function–structure coupling and clinical variables in SZ patients. Specifically, the function–structure differences in WM were extracted, and the SWALFF/FA ratio was calculated as the function–structure coupling to reflect a mismatch between two modality feature alterations (Ji, et al., 2019). Following the normality test by the Kolmogorov-Smirnov test, Pearson’s correlation analyses were subsequently conducted to assess the associations between clinical features (PANSS values and duration of illness) and WM function–structure coupling (SWALFF/FA ratio) in SZ patients.

### 2.7 Associations between WM activation and behaviours

To further examine the relationship between the WM activation and behaviour variables, an independent sample (HCP database, “100 unrelated subjects” dataset release available at http://db.humanconnectome.org.) (Van Essen, et al., 2013), including multimodal MRI data and behaviour data from 100 unrelated healthy subjects, was used in this study. Canonical correlation analyses and Pearson’s correlation analyses were used to estimate the associations between WM activations and five behaviour assessment categories (cognition, emotion, motor, personality and sensory). Detailed information about the behaviour analysis is shown in the Supplementary Material.

### 2.8 Reproducibility analysis on a test–retest database

To investigate the reproducibility and repeatability of SWALFF and dSWALFF in the detection of WM activation, a publicly available test–retest database from the Consortium for Reliability and Reproducibility (CoRR, http://fcon_1000.projects.nitrc.org/indi/CoRR/html/data_citation.html) (Zuo, et al., 2014) was used to retest SWALFF and dSWALFF maps. The NHU database (http://dx.doi.org/10.15387/fcp_indi.corr.hnu1) from the CoRR includes 30 healthy adults (mean age: 24.4 years, range: 20–30 years, 15 female). Each participant received ten scans across one month, one scan occur every three days. This HNU database was used to test the reproducibility of WM activation in different scans. The detailed subject information from the HNU database and methods are shown in the Supplementary Material.

## 3. Results

### 3.1 Subject information

Four SZ patients and four HCs were excluded from this study because they failed to accomplish all of the T1, resting-state fMRI and DTI data acquisitions, or head motion in these scans was larger than 2 mm or 2°. Two groups showed no significant differences in age, gender, education, FD or WM volume (P > 0.05). Information on the subjects is provided in Table 1.

### 3.2 Differences in FA, SWALFF and dSWALFF between SZ patients and HCs

Two-sample t-tests (P<0.05, FWE correction) showed widespread FA reductions within a WM skeleton representing all major WM fasciculi in SZ patients and found a significant increase in FA only in the left posterior limb of the internal capsule (PLIC) in the SZ group (Figure 2A). We also applied another WM microstructural measure of the mean diffusivity (MD). The differences in the MD between SZ patients and HCs showed very high spatial overlap with the results of the FA measure (Figure S1), which further validated the widespread WM microstructural deficits in SZ patients.

**Figure 2.**
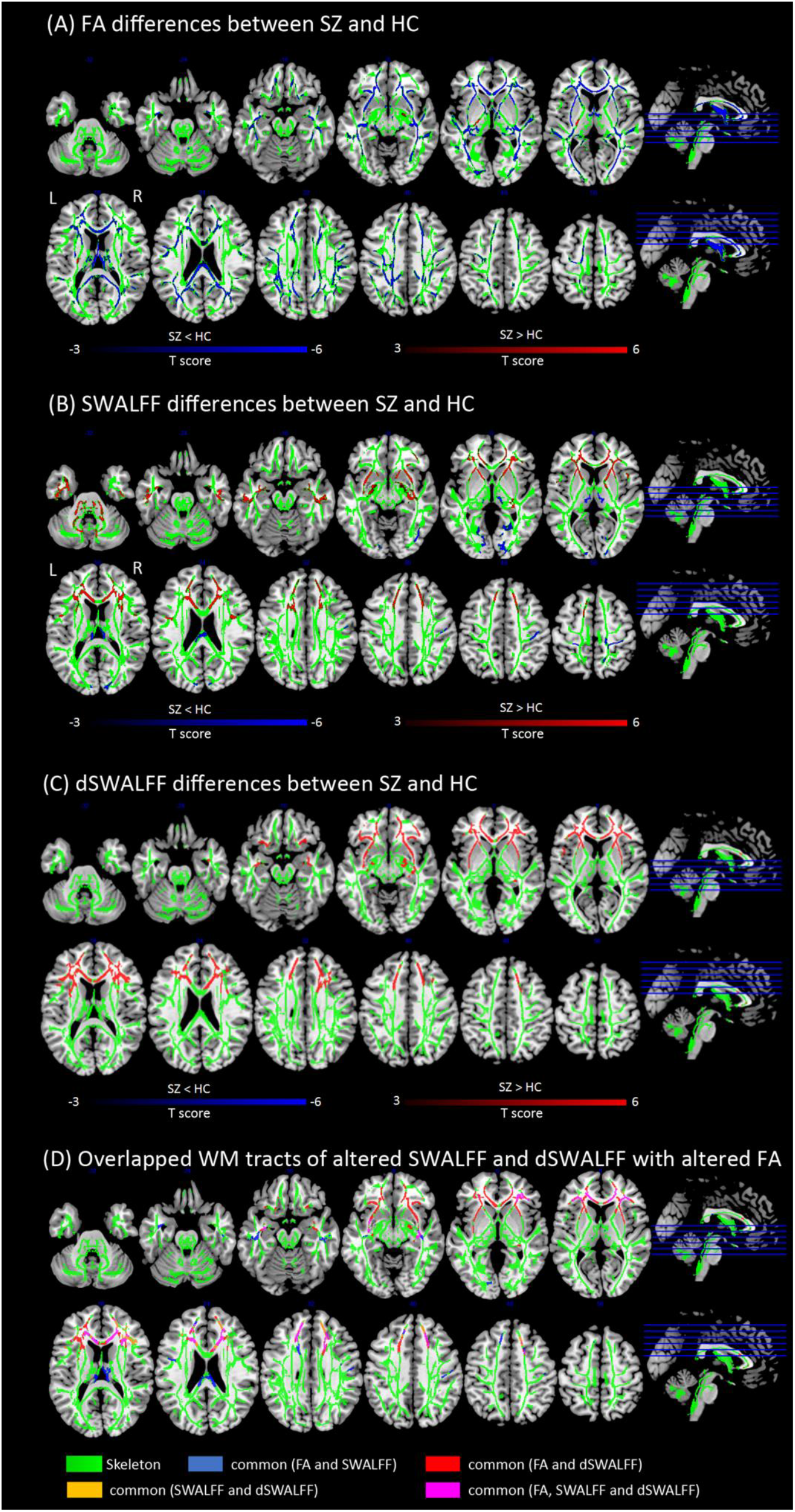
(A) Differences of fractional anisotropy (FA) skeleton between schizophrenia (SZ) and healthy controls (HCs). Two sample T tests (P<0.05, FWE correction) showed widespread FA reductions within a white matter (WM) skeleton representing all major WM fasciculi in SZ, and found a significant increased FA only in the left posterior limb of internal capsule (PLIC). (B) Differences of skeleton-based white matter amplitude of low frequency fluctuation (SWALFF) between SZ and HCs. In SZ, enhanced SWALFF in WM tracts within frontal area (anterior corona radiata [ACR] and genu of corpus callosum [GCC]), temporal regions and anterior limb of internal capsule (ALIC) as well as cerebellum (P<0.05, FWE correction). Decreased SWALFF was observed in WM fibers in occipital area, post-central regions, fornix and splenium of corpus callosum (SCC) compared with HCs (P<0.05, FWE correction). (C) Differences of dynamic SWALFF between SZ and HCs. Increased dynamic SWALFF was observed in WM tracts of frontal area (ACR, GCC and orbitofrontal region [OBF]), external capsule and hippocampus in SZ. (D) Overlapped WM tracts of dysfunction (SWALFF-altered and dSWALFF-altered) with microstructural deficits (FA-altered). The green color represents the WM skeleton. The blue color represents the overlapped tracts between FA-altered and SWALFF-altered. The red color represents the overlapped tracts between FA-altered and dSWALFF-altered. The orange color represents the overlapped tracts between SWALFF-altered and SWALFF-altered. The violet color represents the overlapped tracts among FA-altered, SWALFF-altered and dSWALFF-altered.

Compared with HCs, the SZ group exhibited higher SWALFF in WM tracts within the frontal area (anterior corona radiata [ACR] and genu of the corpus callosum [GCC]), temporal regions and anterior limbs of the internal capsule (ALIC) and cerebellum (P<0.05, FWE correction) (Figure 2B). In addition, decreased SWALFF was observed in WM fibres in the occipital area, post-central regions, fornix and splenium of corpus callosum (SCC) of SZ patients compared with that in HCs (P<0.05, FWE correction) (Figure 2B). In addition, increased dSWALFF was observed in the WM tracts of the frontal area (ACR, GCC and orbitofrontal region [OBF]), external capsule and hippocampus in SZ patients compared with that in HCs (P<0.05, FWE correction) (Figure 2C). The effect size (Cohen’s d) maps of differences in FA, MD, SWALFF and dSWALFF are provided in the Figure S2.

To further examine whether different frequency bands and sliding steps could influence the group differences of LFO in WM, two auxiliary analyses were carried out. We added two additional frequency sub-bands (0.01–0.05 Hz and 0.05–0.10 Hz) to re-calculate the SWALFF maps and re-compared the differences between SZ and HC. We observed that the results from sub-bands were consistent with the results from a full band of 0.01–0.10 Hz (Figure S3). In addition to the sliding steps of 5 TRs in the computation of dSWALFF, multiple sliding steps of 3 TRs, 4 TRs, 6 TRs and 7 TRs were also considered to reproduce the main results (Figure S4).

Another COBRE SZ samples showed consistent differences in the FA, SWALFF and dSWALFF, compared with that from HCs (Figure S5).

### 3.3 Overlapping tracts of FA-altered voxels with SWALFF or dSWALFF alterations

Figure 2D shows the WM tracts overlapping with co-alterations in the microstructure (FA) and activation (SWALFF and dSWALFF). These overlapping tracts were located in the bilateral OBF, ACR, GCC, ALIC, external capsule, SCC, hippocampus and fornix (Figure 2D). Furthermore, within these overlapping tracts, the right ACR (Figure 3A) shows increased SWALFF (T=4.16, p<0.001, effect size=0.549, confidence interval [CI]=[0.067, 0.186]) and decreased FA (T=−3.25, p=0.001, effect size=−0.449, CI=[−0.044, −0.011]) in SZ compared to HCs. Increased SWALFF (T=4.08, p<0.001, effect size=0.540, CI=[0.063, 0.182]) and decreased FA (T=−4.13, p<0.001, effect size=−0.563, CI=[−0.053, −0.019]) was found in the GCC in the SZ group (Figure 3B). Significant negative associations between FA and SWALFF were also observed in the right ACR (r=−0.329, p=0.001) and bilateral GCC (r=−0.362, p<0.001) for SZ patients. Conversely, in the HC group, we only found significant positive correlations between FA and SWALFF in the right ACR (r=0.335, p<0.001). In addition, there were negative correlations between FA and dSWALFF in the left external capsule, right OBF and bilateral GCC in SZ (Table S1). These correlations showed significant group differences between SZ and HC by the non-parametric permutation test (Supplementary Materials).

**Figure 3.**
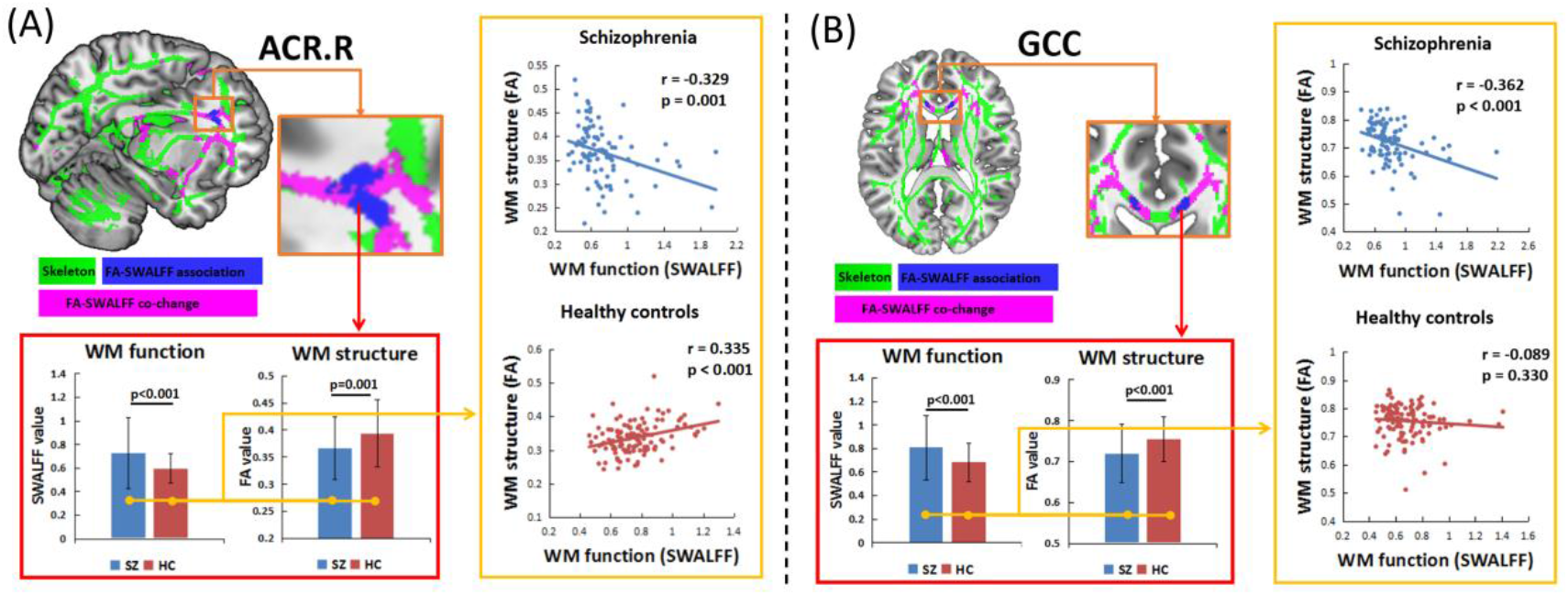
White matter (WM) tract of (A) right anterior corona radiata (ACR) and (B) genu of corpus callosum (GCC) shows increased SWALFF and decreased fractional anisotropy (FA) in schizophrenia (SZ) compared to healthy controls (HCs). The association between FA-altered and SWALFF-altered exhibits the negative correlation in SZ but positive correlation in HCs. Abbreviation: SWALFF, skeleton-based white matter amplitude of low frequency fluctuation.

### 3.4 Relationship between clinical variables and WM function–structure coupling in SZ

There was a significant positive correlation between illness duration and function–structure coupling in the right OBF, right ACR, right hippocampus and bilateral GCC in the SZ group (Table S2). In addition, the function–structure coupling in the right hippocampus was significant negatively correlated with negative PANSS scores (Table S2). Furthermore, these associations remained significant even after controlling for the effects of education, WM volume and antipsychotics. Finally, we conducted a correlation analysis between head motion parameter and SWALFF and dSWALFF to preclude the possible effect of head motion in our results. There was no significant correlation between the FD and SWALFF or dSWALFF (all P > 0.05), which supported that our results were not dependent upon head motion.

### 3.5 Association between WM activations and behaviour scores

Canonical correlation analysis illustrated significant associations between the WM SWALFF and the cognition (R^2^=0.980, P<0.001) as well as emotion (R^2^=0. 973, P<0.001) (Figure 4A1). Additionally, the dSWALFF was significantly associated with the cognition (R^2^=0.909, P=0.003) (Figure 4A2). In addition, Pearson’s correlation analysis showed that the SWALFF in GCC was significantly negatively associated with cognition instruments, including reading decoding, self-regulation/impulsivity and cognition crystallized composite (Ps<0.05, FDR correction) (Figure 4B1). The dSWALFF in the GCC was negatively correlated with self-regulation/impulsivity (P<0.05, FDR correction) (Figure 4B2).

**Figure 4.**
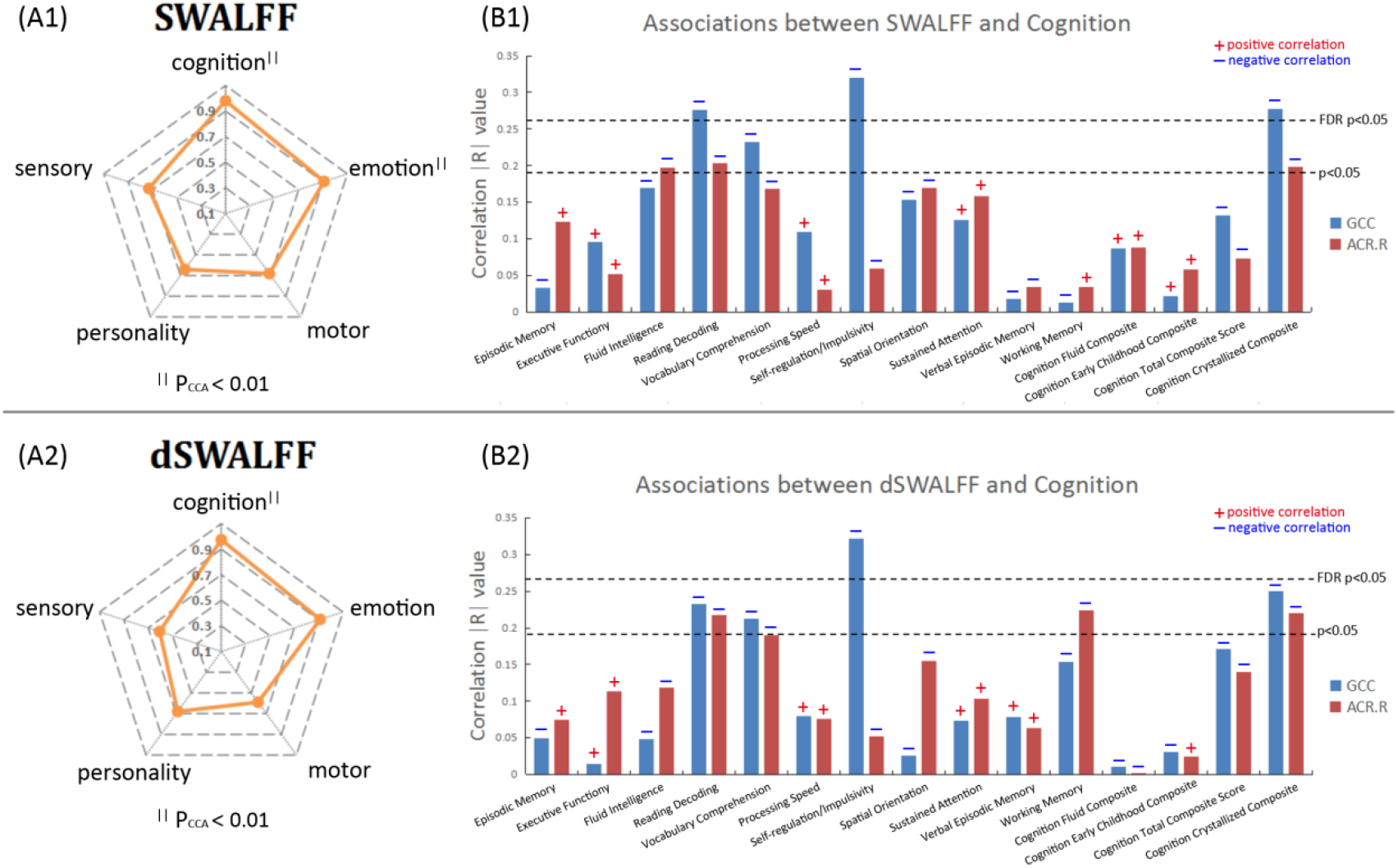
Associations between WM activations and behavior scores. (A) The canonical correlation coefficient between WM activations (SWALFF [A1] and dSWALFF [A2]) and five behavior assessment category (cognition, emotion, motor, personality and sensory). ^∥^ represents a significant positive correlation by the canonical correlation analysis (P_CCA_<0.01). (B) The Pearson’s correlation coefficient between WM activations (SWALFF [B1] and dSWALFF [B1]) in GCC and ACR.R and each cognition instrument.

### 3.6 Reproducibility and repeatability of WM activation

Figure S6 showed the mean correlation coefficient between the SWALFF maps obtained from repeated scans and paired subjects in one scan. The mean coefficients (intra-subject) between the SWALFF maps generated from repeated scans were very high (>0.85). The mean coefficients (inter-subject) between subjects also showed good consistency (>0.65). There were no significant differences between correlations of different paired scans. In total, there was a high degree (>0.8) of consistency within subjects and good consistency (>0.6) between subjects for the SWALFF map. In addition, Figure S7 showed high consistency in SWALFF for WM tracts.

## 4. Discussion

In this study, DTI revealed widespread FA reductions within major WM fasciculi in SZ patients. Meanwhile, using resting-state fMRI, the investigation of WM function showed hyper-activation in WM tracts of the frontotemporal–thalamic–cerebellar pathway and the hypo-activation of primary perception–motor and interhemispheric tracts in SZ patients. We integrated these findings into the global WM abnormalities from both microstructure deficits and dysfunctions, which extended the assumption of the dysconnectivity hypothesis of SZ. The predominant finding of this study indicated a reversed pattern of microstructural deficits and dysfunction in localized WM regions (frontotemporal tracts including ACR, OBF, hippocampus, ALIC and GCC) in SZ patients. Concretely speaking, the key results show that (1) WM activation was significantly increased in SZ patients, while the microstructure FA of WM was reduced; (2) the association between function and structure was significantly positively correlated in HCs but was negatively correlated in SZ patients; and (3) by using the function/structure ratio to measure the coupling of function–structure, we observed that the function/structure ratio was positively related to illness duration and clinical symptoms in SZ patients. Taken together, function–structure association may be one of the potential mechanisms of the dysconnectivity hypothesis of SZ.

### WM global abnormalities: widespread FA decreases

Highly consistent with a prior meta-analysis evaluating DTI data from 1963 SZ patients and 2359 HCs (Kelly, et al., 2018), we found widespread FA reductions within major WM fasciculi in SZ patients. These major fibres included the WM tracts of frontotemporal, occipital, interhemispheric (corpus callosum and fornix) and corticothalamic (corona radiata and ALIC) regions, suggesting widespread WM microstructure deficits in SZ (Kelly, et al., 2018). Reduced FA in these regions has been reported in first-episode and chronic SZ patients and in high-risk individuals (Aydin, et al., 2008; Bora, et al., 2011) and has been associated with illness duration and the severity of positive, negative and cognitive symptoms (Rosenberger, et al., 2012; Whitford, et al., 2010). In addition, this study also observed increased FA in the PLIC in SZ patients, probably because of aberrant axonal pruning and/or fibre organization (Klauser, et al., 2017; Kubicki, et al., 2005). In total, the widespread distribution of FA reductions may support the concept of SZ as a disorder of global structural dysconnectivity.

### WM global abnormalities: LFO hyper-activation and hypo-activation

In addition to the conventional measure (i.e., FA) in WM microstructure, we adopted a novel measure for investigating WM dysfunction in SZ by fMRI. WM dysfunction has been examined in brain disorders such as mild cognitive impairment, epilepsy, Alzheimer’s and Parkinson’s diseases using resting-state fMRI (Chen, et al., 2017; Ji, et al., 2019; Makedonov, et al., 2016). For instance, Jiang and colleagues reported increased functional connectivity within the WM regions located under the Rolandic and precentral/postcentral areas in un-medicated patients with benign epilepsy with centrotemporal spikes (Jiang, et al., 2019b). In the current study, by mapping the resting-state LFO on the WM skeleton, we found widespread LFO abnormalities across WM major tracts, with hyper-activation in frontotemporal regions, ALIC and cerebellum and hypo-activation in occipital and post-central regions compared with activation in HCs. As the ALIC connects the thalamus and frontal cortex, the WM tracts showing hyper-activation may be integrated into the frontotemporal–thalamic–cerebellar pathway. In addition, the occipital and post-central regions are important hubs of visual and sensorimotor processing; thus, the WM pathway showing hypo-activation may be integrated into primary perception–motor tracts. The hyper-activation (hypo-activation) of the frontotemporal–thalamic–cerebellar (primary perception–motor) pathway was consistent with results from previous grey matter LFO studies in SZ (Xu, et al., 2015), indicating multiple impairments, including sensory integration, motor regulation, and cognitive and emotional dysfunction in SZ (Xu, et al., 2015). Furthermore, our findings showed the associations between the resting-state LFO of WM and assessments in cognition and emotion. A negative correlation was demonstrated between the cognitive impulsivity (Barch, et al., 2013) and SWALFF as well as dSWALFF in the GCC. Impulsiveness is a prominent clinical symptom in SZ (Reddy, et al., 2014). Previous studies found that DTI measured WM integrity reduction in frontal regions was associated with increased impulsivity in SZ (Hoptman, et al., 2002). Additionally, this study also found hypo-activation in interhemispheric WM tracts (fornix and SCC), extending the abnormalities of interhemispheric functional communication in SZ (Agcaoglu, et al., 2018; Wang, et al., 2019). In general, these abnormalities in WM activation in the resting state may extend the understanding of the pathophysiology of SZ from the perspective of WM dysfunction.

### Relationship between the function and structure in WM of SZ

The predominant finding of this study indicated a reversed pattern of microstructural deficits and dysfunction in localized WM regions of the ACR, OBF, hippocampus, ALIC and GCC, which can be concluded to be frontotemporal tracts. Concretely speaking, the FA of the frontotemporal tracts was significantly reduced in SZ patients, indicating an impaired structural basis for information transmission (Kelly, et al., 2018). In contrast, the activation of frontotemporal tracts was increased in SZ patients may relate to high functional loading induced by aberrant self-referential processing in SZ (Kuhn and Gallinat, 2013). These results are consistent with previous studies reporting that reduced structural integrity was paralleled by higher functional connectivity in SZ, which may suggest that the impact of structural deficits can be compensated by an increase functional integration (Cocchi, et al., 2014).

More importantly, the functional–structural relationship showed a significant positive correlation in HCs but exhibited a negative correlation in SZ patients. Wu and colleagues have reported that resting-state functional connectivity between white and gray matter shared notable similarities with diffusion- and histology-derived anatomic connectivities in non-human primate, suggesting a positive association between WM function and structure (Wu, et al., 2019). Consistent with Wu’s study, the current study also observed a positive correlation in function–structure coupling in healthy human brain. However, this function–structure relationship was reversed in SZ patients, which may suggest the discordance between higher functional activation and reduced structural integration. In total, these findings suggested the WM function–structure relationship in SZ patients, which may become a possible fundamental mechanism of the dysconnectivity hypothesis of SZ.

### WM function–structure dys-coupling: relationship with clinical variables

In further work, we used the function/structure ratio to measure the coupling of the WM function structure in SZ patients. A higher function/structure ratio reflects more severe dys-coupling. The current study found that the duration of illness was positively correlated with the dys-coupling of function and structure in the ACR, OBF, hippocampus and GCC. Our results were consistent with previous findings showing that the duration of illness was negatively associated with FA (Kelly, et al., 2018) because of the calculation of dys-coupling using the reciprocal value of FA. These associations remained significant even when controlling for education, WM volume and drug effects. This finding is also consistent with many DTI studies reporting no detectable relationship between medication and FA (Kelly, et al., 2018; Samartzis, et al., 2014). As age was highly correlated with illness duration, it was understandable that these associations were no longer significant after controlling for age. In the future, a longitudinal design will be necessary to distinguish the effects of age and illness duration. In addition, the dys-coupling of function–structure in the HIP was positively correlated with negative symptoms in SZ patients, consistent with previous findings of the negative relationships between negative symptoms and FA in WM (Kelly, et al., 2018). Recently, non-invasive neuro-modulation method, such as transcranial magnetic stimulation (TMS), has shown potential to treat SZ (Chen, et al., 2019). WM function may be helpful for understanding how TMS affects the distant cerebral cortex via WM tracts. Taken together, these findings confirmed a positive relationship between the function–structure dys-coupling and the severity of illness in SZ patients.

### Skeleton-based White matter Functional Analysis

Regarding the methodological aspect, fMRI processing in WM should be very careful because WM signals have weaker signal noise ratio than GM and easily mixed by neighboring GM signals due to partial volume effects (Gawryluk, et al., 2014b). Previous studies usually segmented the functional images into GM and WM mask before spatial normalization and smooth (Ji, et al., 2019; Peer, et al., 2017). However, it is very difficult to precisely delineate the optimal WM mask, especially in functional images. Even we determines the optimal WM mask in each subject’s image (i.e., original space), the mask would be non-linearly transformed during spatial normalization. In addition, how to estimate whether the voxel belongs to WM or GM, it should be probabilistic rather than absolute (0 or 1); thus, it may be arbitrary to choose the absolute threshold to make WM mask. Alternatively, we used the DTI (i.e., FA images) to estimate the weighted parameters to those voxels who are closer to the center of the WM tract. Finally, measurement in reliability is required for both research and clinical use (Zuo, et al., 2019). There was a high degree (>0.8) of consistency within subjects and good consistency (>0.6) between subjects for the SWALFF maps from multiple scans in a short time-frame (ten scans across one month), which suggested the reliability of the WM activations. In addition, the WM activation changes were consistent in two independent SZ sample, indicating the sensibility in detection of the disease-related alterations.

### Limitations

The current study has certain limitations. First, most of the SZ participants were chronic and were taking antipsychotic medications. Although no correlation was observed between medication and WM differences, it is difficult to rule out the effect of medication on WM alterations. Second, the present study is a cross-sectional design. As SZ is a progressive psychiatric disorder, a longitudinal design is needed to investigate the dynamic changes in WM dysfunction and microstructural properties in SZ. Third, accounting for cognitive, environmental and genetic factors linked to SZ (Devor, et al., 2017; Van Rheenen, et al., 2018), neuropsychological assessment and genetic research are required for a deeper understanding of WM dysfunctions observed in this study. Finally, previous studies have suggested that the hemodynamic alterations in the BOLD signal in WM may be associated with the energy demands for spiking activity, the maintenance of resting potential and housekeeping processes (Gawryluk, et al., 2014b). However, unlike grey matter, the physiological basis of functional activation in WM remains undetermined. In addition, future work should further address some methodological issues, such as the effects of head motion, respiration and and other artefacts on WM function.

### Conclusion

This study indicated global WM abnormalities and a reversed pattern of microstructural deficits and dysfunction in localized WM tracts in frontotemporal regions in SZ patients, which suggested function–structure dys-coupling with increased functional activations and reduced structural integration in SZ patients. Furthermore, function–structure dys-coupling was positively related to illness duration and clinical symptoms in patients with SZ. In conclusion, function–structure dys-coupling may be one of the potential mechanisms of the dysconnectivity hypothesis of SZ.

## Acknowledgements

The HCP database were provided by the Human Connectome Project, WU-Minn Consortium (Principal Investigators: David Van Essen and Kamil Ugurbil; 1U54MH091657) funded by the 16 NIH Institutes and Centers that support the NIH Blueprint for Neuroscience Research; and by the McDonnell Center for Systems Neuroscience at Washington University.

The COBRE database was downloaded from the COllaborative Informatics and Neuroimaging Suite Data Exchange tool (COINS; http://coins.mrn.org/dx) and data collection was performed at the Mind Research Network, and funded by a Center of Biomedical Research Excellence (COBRE) grant 5P20RR021938/P20GM103472 from the NIH to Dr.Vince Calhoun.

The CoRR NHU database was downloaded from the Consortium for Reliability and Reproducibility (http://fcon_1000.projects.nitrc.org/indi/CoRR/html/data_citation.html) (Principal Investigators: Xu-Chu Weng and Xi-Nian Zuo) and data collection (Thanks for Bing Chen and Qiu Ge) was performed at the Center for Cognition and Brain Disorders, Hangzhou Normal University, Gongshu, Hangzhou, Zhejiang 310036, China, and funded by National Natural Science Foundation of China (31070905, 31371134) and National Social Science Foundation of China (11AZD119).

## Contributors

Cheng Luo and Dezhong Yao designed the study and supervised the project; Shicai Li, Xufeng Song and Mingjun Duan managed the experiments and data collection; Yuchao Jiang, Xiangkui Li, Huan Huang and Xuan Li undertook the data analysis; Yuchao Jiang wrote the manuscript. All authors reviewed the manuscript and approved the final manuscript.

## Data availability

The data supporting the findings of this study are available from the corresponding author upon reasonable request.

