## Supplementary Materials for "Function-structure Coupling: White matter fMRI hyper-activation associates with structural integrity reductions in schizophrenia"

**Jiang et al.**

***This supplementary material includes as follows:***

**PART 1: Methods and Materials**

**Methods and Materials:** Subjects

**Methods and Materials:** Data acquisition

**Methods and Materials:** Subject information of the COBRE database

**Methods and Materials:** Behaviour analysis on the HCP database

**Methods and Materials:** Non-parametric permutation tests

**Methods and Materials:** Reproducibility analysis on a test-retest database (CoRR-NHU)

**PART 2: Supplementary Tables (Table S1-S2)**

**PART 3: Supplementary Figures (Figure S1-S7)**

### Methods and Materials: *Subjects*

The current study included ninety-seven schizophrenia (SZ) patients and 126 healthy subjects (HC) from the Clinical Hospital of Chengdu Brain Science Institute (CHCBSI), China. Some of the patients were part of our previous studies and have been listed in a prior published study (Dong, et al., 2019). In addition to this, forty-seven patients were included in a prior study in which the author investigated the static and dynamic functional connectivity between the cerebellum and cortical/subcortical networks (He, et al., 2019). Fifty-four patients were used in a prior study that examined the thalamic structural and functional connections based on diffusion MRI and resting-state functional MRI (Gong, et al., 2019). The same patients dataset have been used in recent studies. Jiang et al. (2020) examined the global topological disruptions of large-scale white matter and gray matter networks using graph theoretical approaches. Different from prior articles, the current study investigated the coupling of the fractional anisotropy and low-frequency oscillation activation in white matter using skeleton-based white matter functional analysis.

### Methods and Materials: *Data acquisition*

High-resolution T1-weighted images, resting-state fMRI and diffusion-weighted images were acquired in a 3.0 Tesla GE MRI scanner (DISCOVERY MR 750, USA) at the Center for Information in Medicine of University of Electronic Science and Technology of China. Resting-state fMRI data were collected based on a standard T2-weighted EPI sequence (repetition time, 2.0 s; echo time, 30 ms; field of view, 240 × 240 $mm^2$; matrix size, 64 × 64; flip angle, 90°; slice thickness, 4 mm with no gap; 35 slices). Totally, 255 volumes were obtained in 510s. During resting-state fMRI scanning, subjects were instructed to keep their eyes closed, remain awake and not fall asleep. High-resolution T1-weighted data were acquired by a 3D FSPGR sequence (repetition time, 6.008 ms; echo time, 1.984 ms; field of view, 256 × 256 $mm^2$; flip angle, 9°; slice thickness, 1 mm with no gap). Diffusion-weighted brain images were collected from 64 directions by using a diffusion-weighted spinecho EPI sequence (b value, 1000 $s/mm^2$; repetition time, 8500 ms; matrix size, 128 × 128; field of view, 25.6 × 25.6 $cm^2$; slice thickness, 2 mm and 78 slices. In addition, three un-weighted images (b0 images) were obtained in the anterior/posterior frequency direction for correction of susceptibility-induced distortions.

### Methods and Materials: *Subject information of the COBRE database*

The Center for Biomedical Research Excellence in Brain Function and Mental Illness (COBRE) has published raw anatomical, diffusion and functional MR data from 62 patients with schizophrenia (ages=38.8±12.9; 13 female) and 71 healthy controls (ages=37.6±12.9; 17 female). Diagnostic information was collected using the Structured Clinical Interview used for DSM Disorders (SCID). For anatomical imaging a multi-echo MPRAGE (MEMPR) sequence was used with the following parameters: TR/TE/TI=2530/[1.64, 3.5, 5.36, 7.22, 9.08]/900 ms, flip angle=7°, FOV=256×256 mm, thickness=176 mm, matrix=256×256×176, voxel size=1×1×1 mm, number of echoes=5, pixel bandwidth=650 Hz, total scan time=6 min. With 5 echoes, the TR, TI and time to encode partitions for the MEMPR are similar to that of a conventional MPRAGE, resulting in similar GM/WM/CSF contrast. Resting state functional MRI (rs-fMRI) data was collected with single-shot full k-space echo-planar imaging (EPI) with ramp sampling correction using the intercomissural line (AC-PC) as a reference (TR: 2 s, TE: 29 ms, matrix size: 64×64, 32 slices, voxel size: 3×3×4mm$^3$). Anatomical MRI, diffusion MRI, resting fMRI and phenotypic data including: gender, age, handedness and diagnostic information, are released for every participant.

### Methods and Materials: *behaviour analysis on the HCP database*

To further examine the relationship between the WM activation and behaviour variables, an independent sample (HCP database, "100 unrelated subjects" dataset release available at http://db.humanconnectome.org.)[1] including multimodal MRI data and behaviour data from 100 unrelated healthy subjects was used in this study.

Multimodal MRI data, including T1-weighted, fMRI and diffusion-weighted imaging were collected from all subjects on a customized Siemens 3T Connectome Skyra scanner using HCP's acquisition protocol [2]. Structural images were acquired using a 3D MPRAGE T1-weighted sequence with 0.7 mm isotropic resolution. Other parameter settings included: TR=2400ms; TE=2.14ms; TI=1000ms; flip angle=8. Two resting-state fMRI sessions were collected for each subject. In each session, two runs were acquired using single-shot EPI with alternating (left-to-right, LR and right-to-left, RL) phase encoding directions. The two resting sessions were acquired on separate days with the following scanning parameters: TR=720ms; TE=33.1ms; flip angle=52; slice thickness=2.0mm; 72 slices; 2 mm isotropic voxels; multiband factor=8; matrix size=104×90; partial Fourier=6/8; echo spacing=0.58 ms; bandwidth (BW)=2290 Hz/px; time points=1200. Full details about subject recruitment and MRI data acquisition can be found in [3, 4].

Behavioral and other individual subject measure data (both NIH Toolbox and non-Toolbox measures) is available on all subjects. In this study, five major categories included cognition, emotion, motor, personality and sensory were considered. The cognition category includes 12 instruments (Episodic Memory (Picture Sequence Memory), Executive Function/Cognitive Flexibility (Dimensional Change Card Sort), Executive Function/Inhibition (Flanker Task), Fluid Intelligence (Penn Progressive Matrices), Language/Reading Decoding (Oral Reading Recognition), Language/Vocabulary Comprehension (Picture Vocabulary), Processing Speed (Pattern Completion Processing Speed), Self-regulation/Impulsivity (Delay Discounting), Spatial Orientation (Variable Short Penn Line Orientation Test), Sustained Attention (Short Penn Continuous Performance Test), Verbal Episodic Memory (Penn Word Memory Test) and Working Memory (List Sorting)). The emotion category includes 5 instruments (Emotion Recognition (Penn Emotion Recognition Test), Negative Affect, Psychological Well-being, Social Relationships and Stress and Self Efficacy). The motor category includes 4 instruments (Endurance (2 minute walk test), Locomotion (4-meter walk test), Dexterity (9-hole Pegboard) and Strength (Grip Strength Dynamometry)). The personality category includes 1 instrument (Five Factor Model (NEO-FFI)). The sensory category includes 6 instruments (Audition (Words in Noise), Olfaction (Odor Identification Test), Pain (Pain Intensity and Interference Surveys), Taste (Taste Intensity Test), Vision (EVA Scores and Farnsworth Test) and Contrast Sensitivity (Mars Contrast Sensitivity)). Full details about behavioral data can be available in (https://wiki.humanconnectome.org/display/PublicData/).

As the HCP fMRI data had a different TR with other database in this study (HCP: TR=0.72s; other database: TR=2s), the HCP fMRI time series were down-sampled into the TR=2.1 second. To further reduce the computation complex, only the first 630 seconds time series were used to analyze. The SWALFF and dSWALFF was calculated using the same procedures as mentioned in the main text. Subsequently, the JHU atlas [5] was used to define 48 WM tracts, which has been widely used to investigate WM activation in previous studies(See following table) [6]. Averaged SWALFF and dSWALFF values were extracted from each WM region to reflect the WM activations. Ten canonical correlation analyses were used to estimate the associations between WM activations (48 SWALFF values and dSWALFF values) and five behaviour assessment category (cognition, emotion, motor, personality and sensory), separately. Final, the SWALFF and dSWALFF of two important WM tracts (GCC and ACR.R) were extracted. Pearson's correlation analyses were used to examine the relationship between WM activation in GCC and ACR.R and each cognition instrument.


***White matter tracts in the JHU atlas***

| JHU number | Full name | Abbreviation | x | y | z |
|---|---|---|---|---|---|
| 1 | Middle cerebellar peduncle | mCBLP | 0 | -42 | -36 |
| 2 | Pontine crossing tract | PC | 0 | -30 | -33 |
| 3 | Genu of corpus callosum | GCC | 0 | 27 | 6 |
| 4 | Body of corpus callosum | BCC | 0 | -6 | 27 |
| 5 | Splenium of corpus callosum | SCC | 3 | -42 | 18 |
| 6 | Fornix | FX | 0 | -6 | 12 |
| 7 | Corticospinal tract R | CST.R | 9 | -24 | -33 |
| 8 | Corticospinal tract L | CST.L | -6 | -24 | -33 |
| 9 | Medial lemniscus R | ML.R | 6 | -36 | -33 |
| 10 | Medial lemniscus L | ML.L | -3 | -36 | -33 |
| 11 | Inferior cerebellar peduncle R | iCBLP.R | 9 | -45 | -39 |
| 12 | Inferior cerebellar peduncle L | iCBLP.L | -9 | -45 | -39 |
| 13 | Superior cerebellar peduncle R | sCBLP.R | 6 | -42 | -24 |
| 14 | Superior cerebellar peduncle L | sCBLP.L | -6 | -42 | -24 |
| 15 | Cerebral peduncle R | CBRP.R | 15 | -18 | -12 |

| | | | | | |
|---|---|---|---|---|---|
| 16 | Cerebral peduncle L | CBRP.L | -12 | -18 | -12 |
| 17 | Anterior limb of internal capsule R | ALIC.R | 18 | 6 | 9 |
| 18 | Anterior limb of internal capsule L | ALIC.L | -18 | 6 | 9 |
| 19 | Posterior limb of internal capsule R | PLIC.R | 21 | -15 | 9 |
| 20 | Posterior limb of internal capsule L | PLIC.L | -18 | -15 | 9 |
| 21 | Retrolenticular part of internal capsule R | RLIC.R | 33 | -30 | 6 |
| 22 | Retrolenticular part of internal capsule L | RLIC.L | -30 | -30 | 6 |
| 23 | Anterior corona radiata R | ACR.R | 21 | 27 | 9 |
| 24 | Anterior corona radiata L | ACR.L | -21 | 27 | 9 |
| 25 | Superior corona radiata R | SCR.R | 24 | -9 | 30 |
| 26 | Superior corona radiata L | SCR.L | -21 | -9 | 30 |
| 27 | Posterior corona radiata R | PCR.R | 24 | -39 | 30 |
| 28 | Posterior corona radiata L | PCR.L | -24 | -39 | 27 |
| 29 | Posterior thalamic radiation R | OR.R | 33 | -57 | 6 |
| 30 | Posterior thalamic radiation L | OR.L | -33 | -57 | 6 |
| 31 | Sagittal stratum R | SS.R | 39 | -33 | -9 |
| 32 | Sagittal stratum L | SS.L | -39 | -30 | -9 |
| 33 | External capsule R | EC.R | 30 | 0 | 3 |
| 34 | External capsule L | EC.L | -30 | 0 | 3 |
| 35 | Cingulum (cingulate gyrus) R | CGG.R | 9 | -9 | 30 |
| 36 | Cingulum (cingulate gyrus) L | CGG.L | -6 | -12 | 30 |
| 37 | Cingulum (hippocampus) R | CGH.R | 24 | -30 | -15 |
| 38 | Cingulum (hippocampus) L | CGH.L | -18 | -33 | -15 |
| 39 | Fornix (cres) / Stria terminalis R | FXC.R | 30 | -24 | -6 |
| 40 | Fornix (cres) / Stria terminalis L | FXC.L | -27 | -24 | -6 |
| 41 | Superior longitudinal fasciculus R | SLF.R | 36 | -27 | 24 |
| 42 | Superior longitudinal fasciculus L | SLF.L | -36 | -27 | 27 |
| 43 | Superior fronto-occipital fasciculus R | SFO.R | 21 | 0 | 21 |
| 44 | Superior fronto-occipital fasciculus L | SFO.L | -21 | 3 | 21 |
| 45 | Uncinate fasciculus R | UF.R | 33 | 0 | -15 |
| 46 | Uncinate fasciculus L | UF.L | -33 | -3 | -15 |
| 47 | Tapetum R | TAP.R | 30 | -45 | 15 |
| 48 | Tapetum L | TAP.L | -27 | -48 | 15 |

### Methods and Materials: *Reproducibility analysis on a test-retest database(CoRR-NHU)*

To investigate the reproducibility and repeatability of SWALFF and dSWALFF in detection of WM activation, a publicly available test-retest database from the Consortium for Reliability and Reproducibility (CoRR, http://fcon_1000. projects.nitrc.org/indi/CoRR/html/data_citation.html) was used. The Consortium for Reliability and Reproducibility (CoRR) is working to address this challenge and establish test-retest reliability as a minimum standard for methods development in functional connectomics. The CoRR has aggregated 1,629 typical individuals' resting state fMRI (rfMRI) data (5,093 rfMRI scans) from 18 international sites, and is openly sharing them via the International Data-sharing Neuroimaging Initiative (INDI). We choose the HNU database because it is a between-sessions repeat experimental design, which is believed to be developmentally stable in a short time-frame. The NHU database (http://dx.doi.org/10.15387/fcp_indi.corr.hnu1) from the CoRR includes 30 healthy adults (mean age: 24.4 years, range: 20-30 years, 15 female). Each participant received ten scans across one month, one scan every three days. Five modalities (EPI/ASL/T1/DTI/T2) of brain images were acquired for all subjects. During functional scanning, subjects were presented with a fixation cross and were instructed to keep their eyes open, relax and move as little as possible while observing the fixation cross. Subjects were also instructed not to engage in breath counting or meditation. More scan parameters can be seen in the website (http://fcon_1000.projects.nitrc.org/indi/CoRR/html/_static/scan_parameters/HNU_1_scan table.pdf).

This HNU database was used to test the reproducibility of the WM activation in different scans. First, the images data were processed by the same procedures as mentioned in the main text and calculated the SWALFF map and dSWALFF map for each scan and each subject. For each subject, we computed the correlation coefficient between any two scans; then, the correlation coefficients were averaged for each scan. Final, the individual-wise correlation coefficients were averaged across all subjects to reflect the group-level intra-subject consistency. In addition, the correlation coefficient between subjects in each scan was also computed to assess the inter-subject consistency. Intra-subject and inter-subject consistency was displayed in Figure S6.

To better illustrate the consistency of SWALFF and dSWALFF across the ten scans, we extracted several important WM tracts to figure the SWALFF and dSWALFF changes across scans. Specifically, the JHU atlas was used to define 48 WM tracts, which has been widely used to investigate WM activation in previous studies. As the cerebellum was not scanning in the HNU database, only 36 WM tracts were considered in this study (See following table). Averaged SWALFF and dSWALFF values were extracted from each WM tract and displayed on Figure S7.

### Methods and Materials: ***non-parametric permutation tests***

To further examine whether the correlations between the FA and SWALFF as well as dSWALFF are significantly different between SZ and HC groups, we conducted a non-parametric permutation test to identify the group interaction. In brief, we calculated the correlation coefficient between FA and SWALFF (dSWALFF) in the SZ and HC group separately. Next, we computed the real difference value of the correlation coefficient between the two groups. Then, we randomly assigned the group labels across all subjects and divided all subjects into two random groups. In each random group, we calculated the correlation coefficient between FA and SWALFF (dSWALFF) and re-calculated the difference in correlation coefficient between the two random groups. Finally, this randomization procedure was repeated 100,000 times, and thus yielded a distribution of the null hypothesis. According the location of the real difference value within distribution of the null hypothesis, a p-value was assigned to the real difference in correlation. The non-parametric permutation test revealed that these correlations showed significant group differences between SZ and HC ($Ps<0.05$).

### Supplementary Tables

**Table S1.** WM Regions with significant associations between FA and dSWALFF in SZ

| Regions | MNI coordinate | | | r-value | P-value | Voxels |
|---|---|---|---|---|---|---|
| | x | y | Z | | | |
| Left EX | -31 | 11 | -2 | -0.351 | 0.00054 | 141 |
| Right GCC | 10 | 30 | -1 | -0.328 | 0.00128 | 58 |
| Left GCC | -8 | 29 | 12 | -0.365 | 0.00034 | 91 |
| Right OBF | -25 | 33 | 1 | -0.379 | 0.00011 | 94 |

Abbreviation: FA, fractional anisotropy; dSWALFF, dynamic skeleton-based white matter amplitude of low frequency fluctuation; WM, white matter; SZ, schizophrenia; EX, external capsule; GCC, genu of corpus callosum; OBF, orbitofrontal.

**Table S2.** Relationships between clinical variables and white matter function-structure coupling in SZ

| Clinical variables | Covariates | r values between clinical variables and WM regions | | | | |
|---|---|---|---|---|---|---|
| | | Right HIP | Right OBF | Left GCC | Right GCC | Right ACR |
| Illness duration | None | **0.464***** | **0.312**** | **0.399***** | **0.381***** | **0.394***** |
| | Education | **0.453***** | **0.264*** | **0.329**** | **0.311*** | **0.307*** |
| | WM volume | **0.476***** | **0.273*** | **0.321*** | **0.353**** | **0.309*** |
| | Antipsychotics | **0.432***** | 0.248 | **0.300*** | **0.301*** | **0.292*** |
| | Age | **0.306*** | 0.165 | **0.342**** | 0.143 | 0.206 |
| PANSS-N | None | **0.253*** | 0.162 | 0.016 | 0.144 | -0.015 |
| | Education | **0.290*** | 0.186 | 0.096 | 0.175 | 0.031 |
| | WM volume | **0.256*** | 0.158 | 0.016 | 0.145 | -0.022 |
| | Antipsychotics | **0.268*** | 0.165 | 0.015 | 0.145 | -0.016 |
| | Age | 0.180 | 0.114 | -0.003 | 0.072 | -0.075 |

Abbreviation: WM, white matter; SZ, schizophrenia; HIP, hippocampus; OBF, orbitofrontal; GCC, genu of corpus callosum; ACR, anterior corona radiata.

The r values represent the associations between clinical variables and WM regions when controlling the effects of covariates.

*** represents the p-value < 0.001

** represents the p-value < 0.01

* represents the p-value < 0.05

**Supplementary Figure S1**

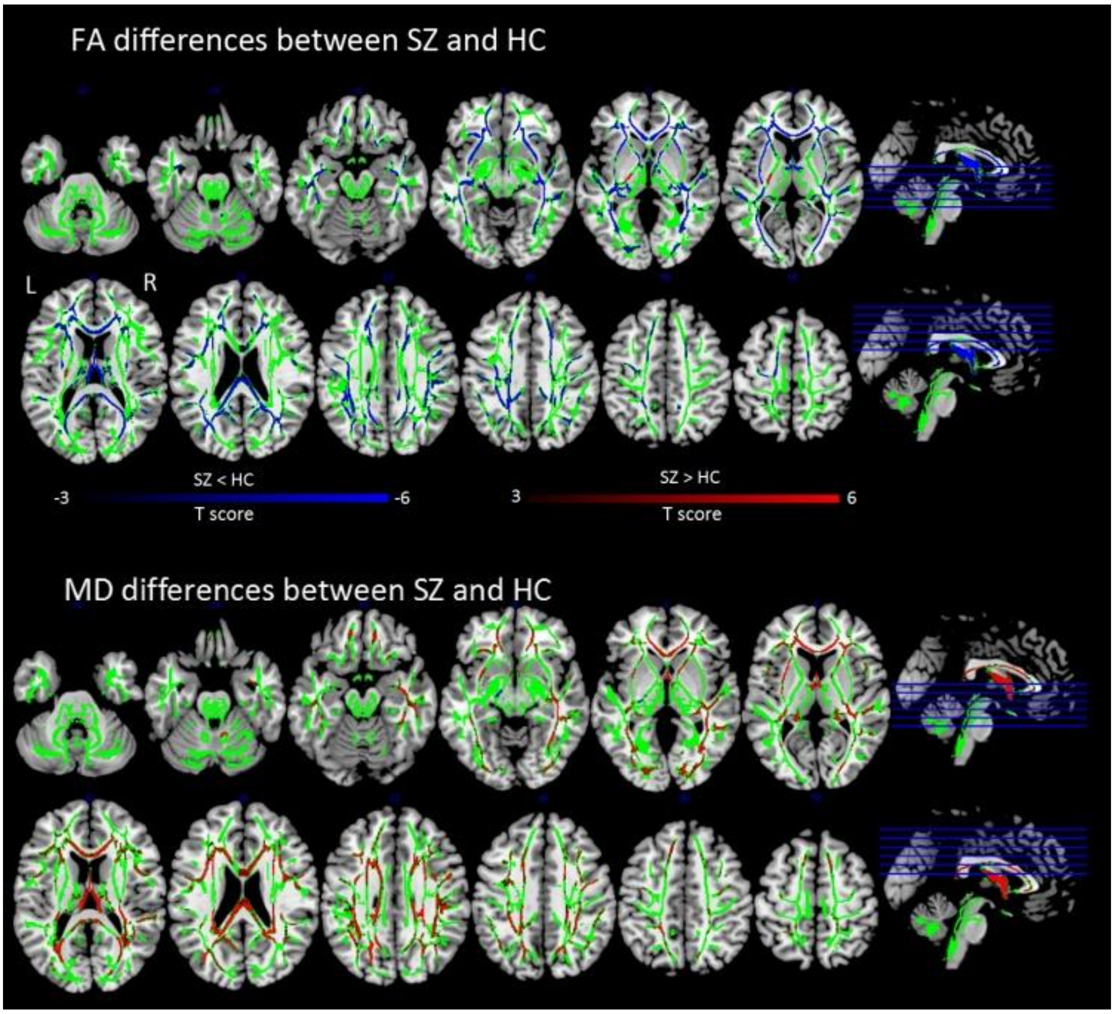


**Figure S1.** Differences of mean diffusivity (MD) between schizophrenia (SZ) and healthy controls (HC) showed very high spatial overlap with the differences between SZ and HC using the FA measure.

### Supplementary Figure S2

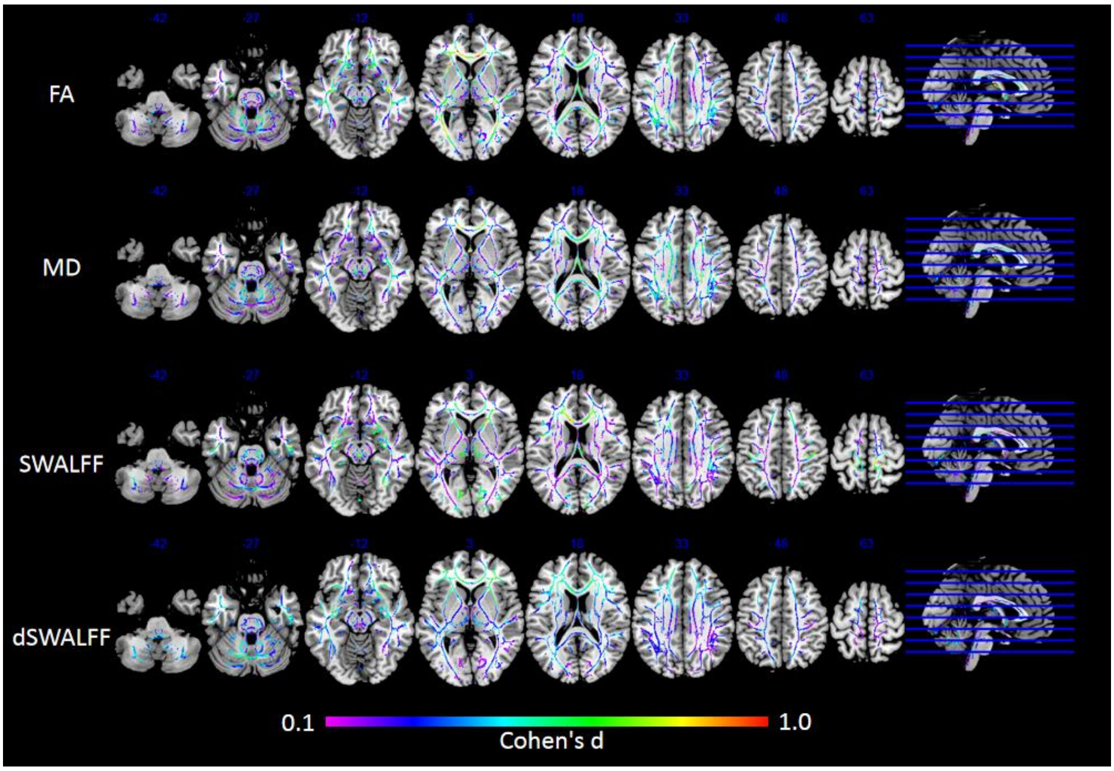


**Figure S2.** Effect size maps of differences in FA, MD, SWALFF and dSWALFF. Effect size was measured by the Cohen's d.

### Supplementary Figure S3

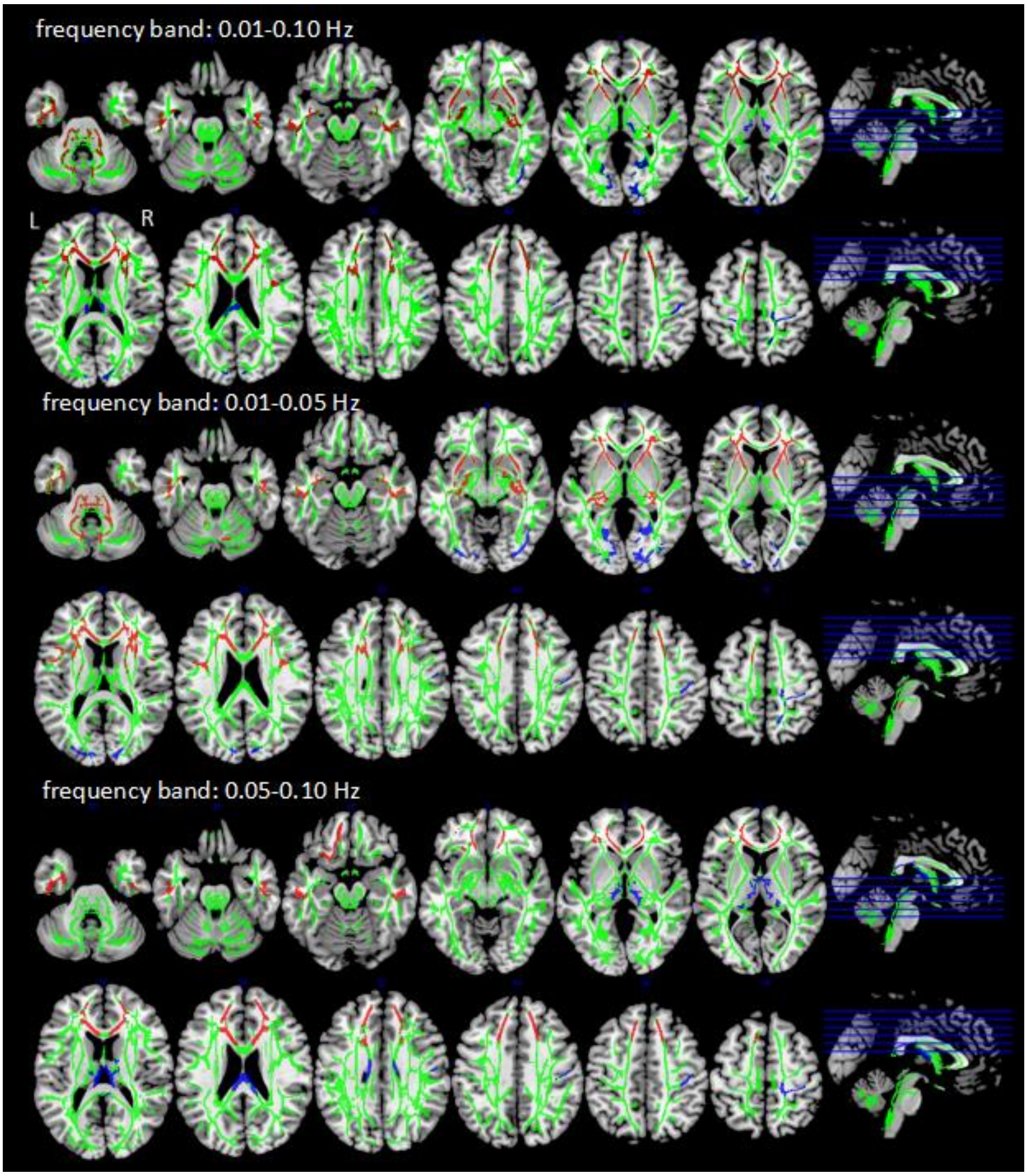


**Figure S3.** Differences of SWALFF between schizophrenia (SZ) and healthy controls (HC) by using the frequency band of 0.01-0.1 Hz, 0.01-0.05 Hz and 0.05-0.10 Hz. The results from frequency sub-band of 0.01-0.1 Hz were consistent with the results from two sub-band of 0.01-0.05 Hz and 0.05-0.10 Hz.

### Supplementary Figure S4

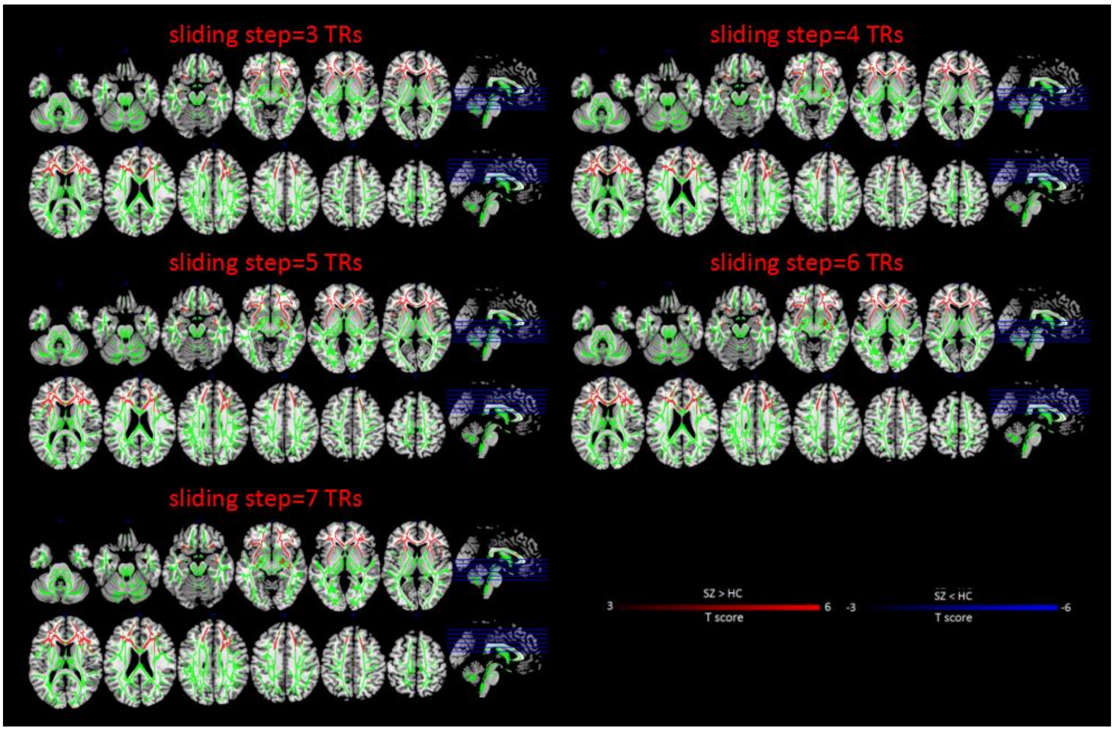


**Figure S4.** Results in differences of dynamic SWALFF between schizophrenia (SZ) and healthy controls (HC) by using different sliding steps (3TRs, 4 TRs, 5 TRs, 6 TRs and 7 TRs).

### Supplementary Figure S5

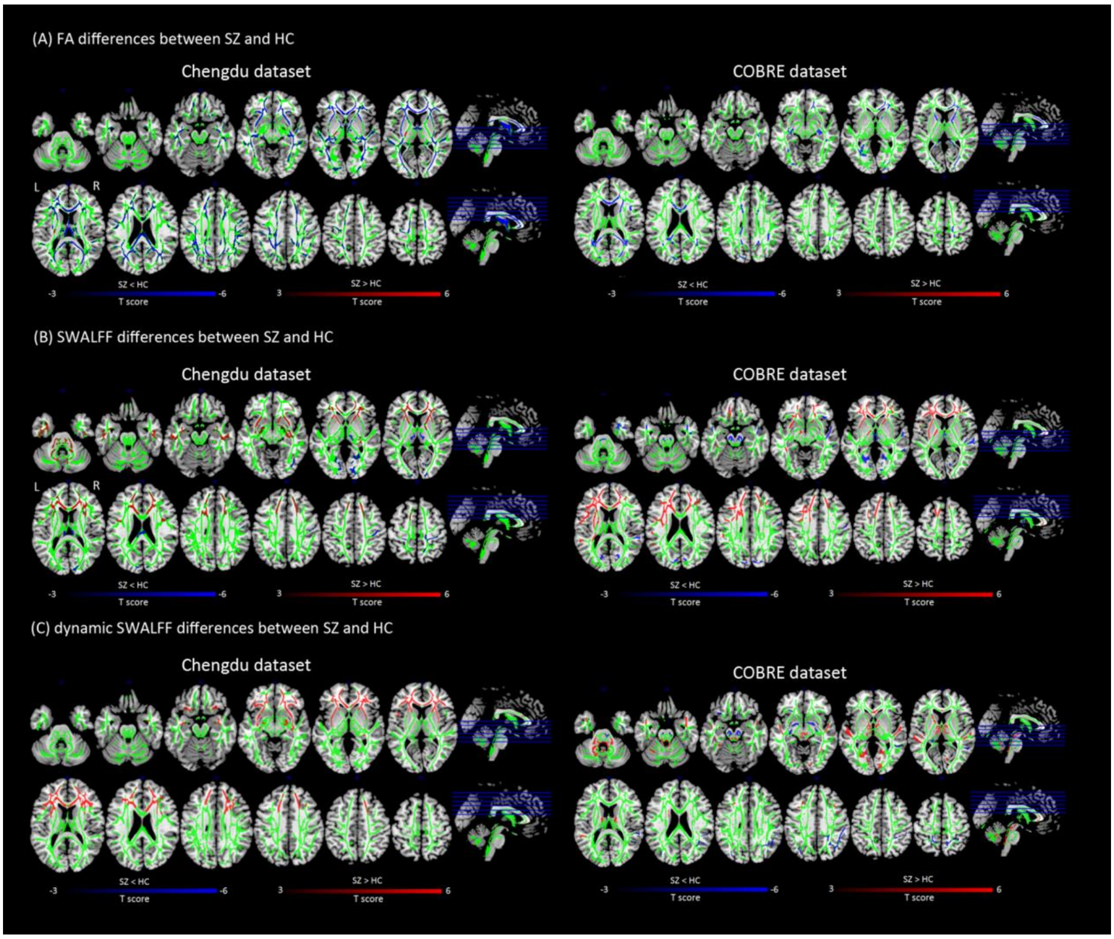


**Figure S5.** Differences of FA, SWALFF and dSWALFF between SZ and HC from two independent samples (Chengdu database and COBRE database). Compared with HC, both two SZ samples showed similar alteration patterns in the white matter skeleton FA, SWALFF and dSWALFF. In particular, both two SZ samples showed widespread decreased FA in corpus callosum, corona radiata and internal capsule. In addition, both two patient samples exhibited increased SWALFF in anterior corona radiata and genu of corpus callosum and reduced SWALFF in occipital area. Both two SZ groups observed increased dSWALFF in anterior corona radiata, corpus callosum and internal capsule. These results illustrated consistent changes in WM microstructure and activation, even in two independent SZ samples.

### Supplementary Figure S6

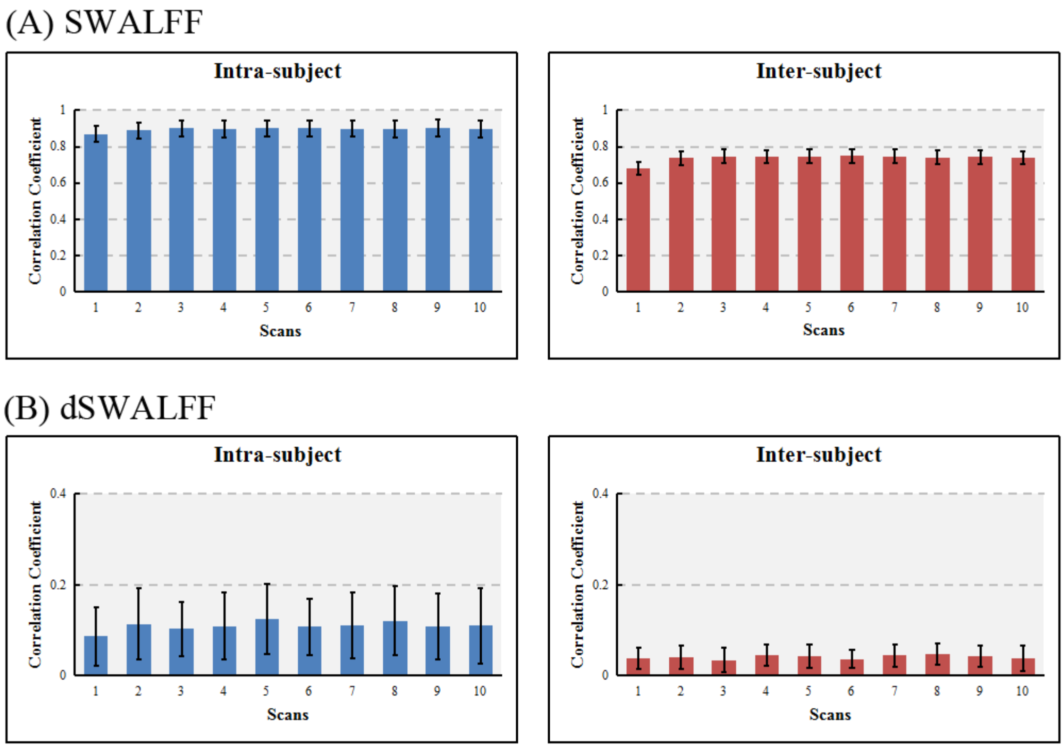


**Figure S6.** Intra- and inter-subject correlation coefficient for (A) SWALFF and (B) dSWALFF. Figure S6 shows the mean correlation coefficient between the SWALFF (dSWALFF) maps obtained from repeated scans and from paired subjects in one scan. The mean coefficients (intra-subject) between the SWALFF maps generated from repeated scans were very high (>0.85). The mean coefficients (inter-subject) between subjects also show a good consistency (>0.65). There were no significant differences between correlations of different paired scans. In total, there is a high degree (>0.8) of consistency within subjects and good consistency (>0.6) between subjects for the SWALFF map. However, correlation coefficients between the dSWALFF maps generated from paired scans or subjects were very low (<0.2). It is understandable because the dSWALFF is a measure of variability rather than stability in brain activity over scanning time.

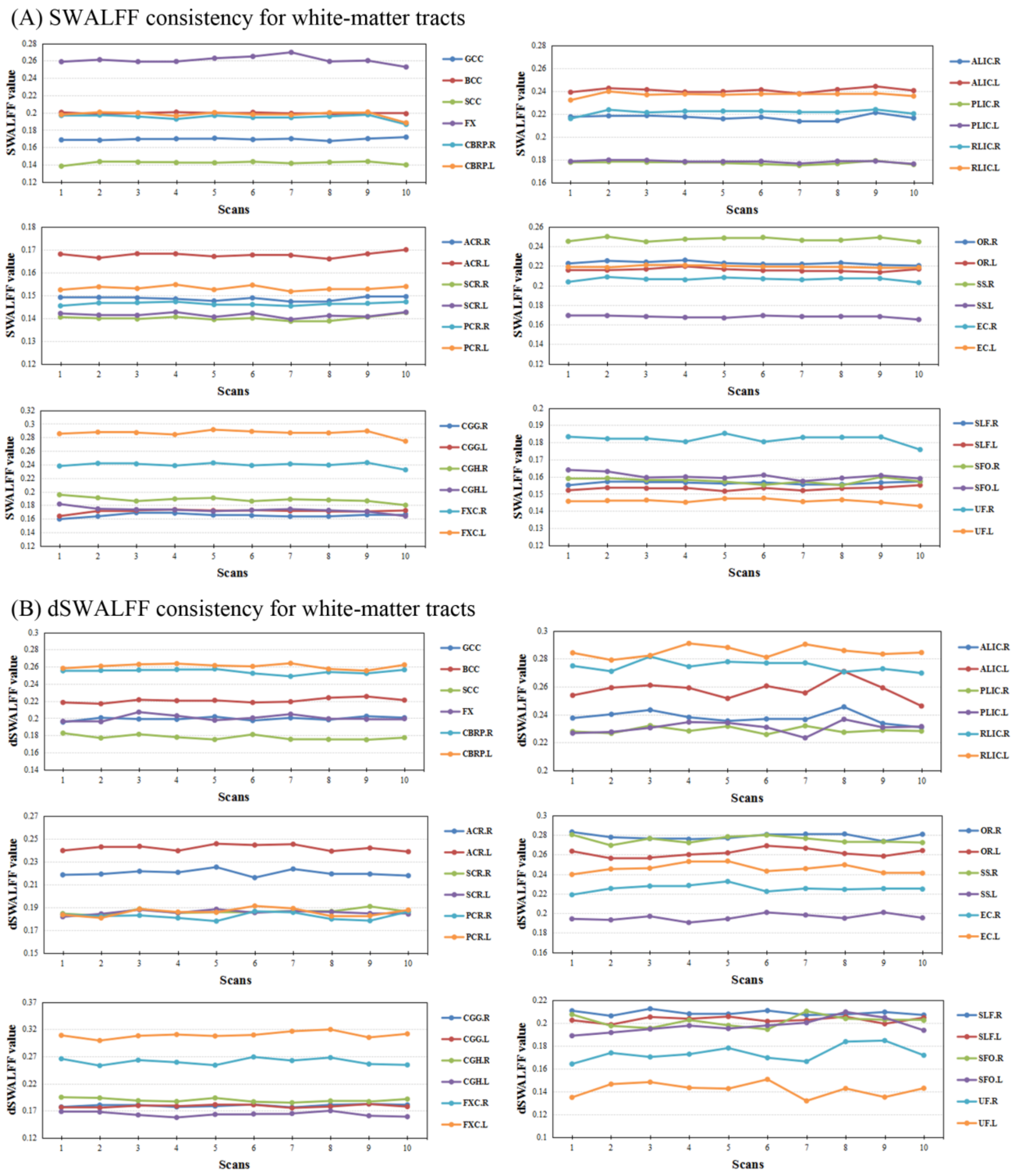


**Figure S7.** The consistency of each WM tract of JHU atlas in (A) SWALFF and (B) dSWALFF at ten scans.
